# Hsp90 promotes kinase evolution

**DOI:** 10.1101/006411

**Authors:** Jennifer Lachowiec, Tzitziki Lemus, Elhanan Borenstein, Christine Queitsch

## Abstract

Heat-shock protein 90 (Hsp90) promotes the maturation and stability of its client proteins, including many kinases. In doing so, Hsp90 may allow its clients to accumulate mutations as previously proposed by the capacitor hypothesis. If true, Hsp90 clients should show increased evolutionary rate compared to non-clients; however, other factors, such as gene expression and protein connectivity, may confound or obscure the chaperone’s putative contribution. Here, we compared the evolutionary rates of many Hsp90 clients and non-clients in the human protein kinase superfamily. We show that Hsp90 client status promotes evolutionary rate independently of, but in a small magnitude similar to that of gene expression and protein connectivity. Hsp90’s effect on kinase evolutionary rate was detected across mammals, specifically relaxing purifying selection. Hsp90 clients also showed increased nucleotide diversity and harbored more damaging variation than non-client kinases across humans. These results are consistent with the central argument of the capacitor hypothesis that interaction with the chaperone allows its clients to harbor genetic variation. Hsp90 client status is thought to be highly dynamic with as few as one amino acid change rendering a protein dependent on the chaperone. Contrary to this expectation, we found that across protein kinase phylogeny Hsp90 client status tends to be gained, maintained, and shared among closely related kinases. We also infer that the ancestral protein kinase was not an Hsp90 client. Taken together, our results suggest that Hsp90 played an important role in shaping the kinase superfamily.

## Introduction

The conserved heat shock protein Hsp90 facilitates the proper folding and stability of its substrates (clients) (Taipale, et al. 2010), many of which are kinases with important roles in growth and development. Hsp90 perturbation increases the penetrance of expressed genetic variants and reveals cryptic genetic variation in genetically divergent populations of plant, fly, yeast, and fish (Rutherford and Lindquist 1998; Queitsch, et al. 2002; Yeyati, et al. 2007; Jarosz and Lindquist 2010). In worms, naturally varying Hsp90 levels predict mutation penetrance with lower Hsp90 levels resulting in greater penetrance (Burga, et al. 2011; Casanueva, et al. 2012). These observations with traditional model organisms prompted the controversial hypothesis that Hsp90 plays an important evolutionary role, allowing genetic variation to remain phenotypically silent and releasing it in environments that perturb Hsp90 function (Rutherford and Lindquist 1998). Consistent with this hypothesis, perturbing Hsp90 function in surface-dwelling *Astyanax mexicanus* fish results in eye phenotypes that are reminiscent of the natural adaptation of eye loss in the cave-dwelling fish of the same species, presumably due to release of Hsp90-dependent standing variation (Rohner, et al. 2013). Hsp90-dependent standing variation occurs frequently in natural strains of plants, flies, and yeast and often affects complex traits (Rutherford and Lindquist 1998; Sangster, Salathia, Lee, et al. 2008; Jarosz and Lindquist 2010), consistent with a significant role of Hsp90 in evolution, especially in the evolution of genes encoding its client proteins.

The evolutionary rate of protein coding genes is commonly measured as the ratio of non-synonymous changes to synonymous changes, dN/dS. Using this measure, we recently reported that genes encoding Hsp90 clients tend to evolve faster than genes encoding their non-client paralogs (Lachowiec, et al. 2013). This trend was not observed without considering paralog status, presumably because many other factors, some of which are shared among paralogs, influence evolutionary rate. One drawback of this study was the small number of available client and non-client paralog pairs. Recently Taipale et al. (2012) systematically annotated Hsp90 clients in the human kinome, using a high-throughput assay to assess kinase and Hsp90 interactions. Of the 314 tested kinases, 98 are classified as strong Hsp90 clients and 95 as weak clients. Notably, the dN of strong Hsp90 clients is greater than the dN of non-clients, suggesting that interaction with Hsp90 may indeed allow for increased accumulation of non-synonymous genetic variation (Taipale, et al. 2012).

There is precedent for chaperone-facilitated evolution of client proteins. In prokaryotes, GroEL/ES clients show different degrees of dependency on the chaperone for stability and folding. Genes that encode proteins with greater GroEL/ES dependency show greater dN/dS after correction for confounding factors, (Fares, et al. 2002; Bogumil and Dagan 2010; Williams and Fares 2010; Bogumil and Dagan 2012; Bogumil, et al. 2012). Similarly, over-expression of GroEL/ES promotes tolerance of non-synonymous mutations (Tokuriki and Tawfik 2009). Factors that may confound putative chaperone effects on evolutionary rate include absolute expression levels (Pál, et al. 2001; Wall, et al. 2005) and a protein’s connectivity in protein-protein interaction (PPI) networks (Fraser, et al. 2002). Highly expressed genes tend to evolve slower than genes with lower expression levels as do genes encoding proteins that interact with many other proteins, presumably due to the selective pressure to maintain all functional interactions. Neither of the studies discussed above (Taipale, et al. 2012; Lachowiec, et al. 2013) controlled for these factors when examining the role of Hsp90 on protein evolution. Another factor contributing to dN/dS is protein stability; genes encoding stable proteins tend to evolve faster (Bloom, et al. 2006). Many Hsp90 clients are inherently unstable and are rapidly degraded in Hsp90-limited conditions (Taipale, et al. 2010). Interaction with the chaperone Hsp90 promotes client protein stability (Taipale, et al. 2012), suggesting a mechanistic basis for the chaperone’s putative effect on dN/dS.

Here we set out to dissect and compare the contributions of Hsp90 client status, gene expression levels and protein interaction degrees to the evolutionary rate of kinases. We evaluate the role of Hsp90 in kinome evolution within and across lineages. Hsp90 client status is thought to be dynamic throughout gene family evolution with as few as one amino acid change resulting in a switch from non-client to client (Citri, et al. 2006; Taipale, et al. 2012; Lachowiec, et al. 2013). We assess this unexplored dynamic by quantifying the transition rates between client and non-client states throughout kinase evolution and show that client status switching is infrequent. We also infer the ancestral state of client status among kinases. Taken together, our results support the controversial hypothesis that Hsp90 plays an important role in the evolutionary processes shaping large gene families.

## Results

### Hsp90 client status contributes to dN/dS

Taipale and co-authors (2012) previously reported that strong Hsp90 kinase clients acquire more non-synonymous mutations than non-client kinases, using pairwise dN values from human and mouse. We first extended these analyses by examining evolutionary rate (pairwise dN/dS between human and mouse, Table 1) for non-, weak, and strong clients based on strength of kinase interaction with Hsp90 as defined previously (Taipale, et al. 2012). Evolutionary rate analysis (dN/dS) is the gold standard for assessing the types of selection potentially acting on proteins. The dN/dS for strong Hsp90 clients was significantly greater than the dN/dS for non-clients (Figure 1a, p = 0.0004, Wilcoxon rank sum test); no significant dN/dS difference was observed between weak clients and non-clients. Both of these findings were consistent with the prior findings observed with dN alone. Second, we found that dN/dS values of non-clients are significantly different from the combined dN/dS values of strong and weak clients (p = 0.01221, Wilcoxon rank sum test), which was not previously addressed (Taipale, et al. 2012). None of the kinases showed dN/dS > 1, which would indicate positive selection. Rather, the Hsp90 client kinases showed relaxed purifying selection compared to non-clients, consistent with previous observations in plants and yeast (Lachowiec et al 2013).

**Figure 1.**
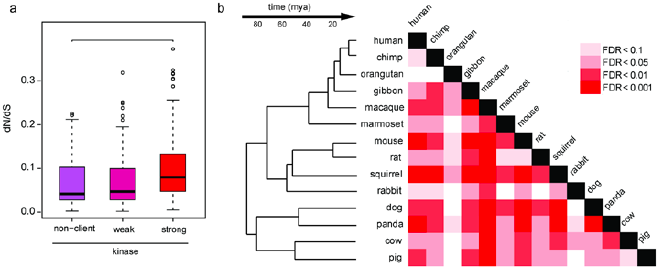
Hsp90 client and non-client kinases differ significantly in evolutionary rate. a) The difference for human-mouse pairwise dN/dS for client and non-client kinases is driven by strong clients (p = 0.0004211, Wilcoxon rank-sum test). b) Significant differences between strong and non-client kinase dN/dS were observed across mammals in a core set of kinases found in all mammalian species examined (Wilcoxon rank-sum test). Shades of red indicate false discovery rate (FDR).

**Table 1.** dN, dS, and dN/dS values for human kinases.

|  | dN | 95% CI for dN | dS | 95% CI for dS | dN/dS | 95% CI for dN/dS |
| --- | --- | --- | --- | --- | --- | --- |
| non-clients | 0.043 | (0.0018, 0.1497) | 0.605 | (0.2704, 1.0417) | 0.069 | (0.0043, 0.2239) |
| all clients | 0.055 | (0.0025, 0.1811) | 0.619 | (0.3106, 1.2571) | 0.088 | (0.0053, 0.3148) |
| weak clients | 0.047 | (0.0011, 0.1780) | 0.629 | (0.2823, 1.2331) | 0.073 | (0.0036, 0.2475) |
| strong clients | 0.063 | (0.0050, 0.1813) | 0.609 | (0.3486, 1.1787) | 0.104 | (0.0091, 0.3176) |

The observed greater dN/dS of Hsp90 clients may be an indirect consequence of other factors that strongly influence evolutionary rate, specifically gene expression levels (Pál, et al. 2001; Wall, et al. 2005) or protein interaction degree (Fraser, et al. 2002). If genes encoding Hsp90 clients are expressed at lower levels (Taipale, et al. 2012) or if client proteins are less connected in protein interaction networks, the correlation between Hsp90 client status and dN/dS can be explained without invoking Hsp90 as a contributor to evolutionary rate. We examined whether gene expression level or PPI connectivity explains the observed contribution of Hsp90 client status to kinase dN/dS by conducting a linear regression analysis. To this end, we calculated the mean and maximum gene expression of client and non-client kinases across RNA-seq experiments for 11 different human primary tissue samples (Castle, et al. 2010) and used PPI values from Taipale et al. (2012). To account for the phylogenetic relatedness among these kinases, we used the kinase tree (Manning, et al. 2002) to calculate phylogenetically independent contrasts (PIC) (Felsenstein 1985) to correct for non-independence of all variables under study: kinase dN/dS, Hsp90 client status, gene expression levels, and PPI values. For this analysis, Hsp90 client status for each kinase was measured by its quantitative interaction score with the chaperone (Hsp90 interaction score, HIS) (Taipale *et al*. 2012). We considered whether expression breadth, measured as the number of tissues in which a kinase is expressed, was correlated with dN/dS. In this data set, however, 187 of the tested 210 kinases were expressed in all tissues, precluding analysis.

We used linear regression to understand the relationships among the PIC for each pair of variables. The PIC of kinase evolutionary rates and Hsp90 interaction score were positively associated (p = 0.0001, Table 2), consistent with our phylogenetically naïve, categorical analysis (Figure 1a). As expected the PIC of kinase evolutionary rate and gene expression levels were negatively associated (p = 0.02, Table 2) (Pál, et al. 2001); the PIC of kinase evolutionary rate and PPI connectivity were also negatively associated (Fraser, et al. 2002) (p = 0.0001, Table 2). Notably, we did not observe a significant association between Hsp90 interaction score and expression contrasts (p = 0.7, Table 2), suggesting that there are no systematic differences in gene expression levels that may drive the observed differences in evolutionary rate between Hsp90 client and non-client kinases. However, we found a negative correlation between PPI connectivity and Hsp90 interaction score contrasts (Table 2), raising the possibility that the greater evolutionary rate of Hsp90 kinase clients derives from fewer protein interactions.

**Table 2.** Hsp90 interaction score is positively associated with dN/dS^1^.

| model | regression coefficient |  | R<sup>2</sup> |
| --- | --- | --- | --- |
| dN/dS ~ Hsp90 interaction score | 0.007462 | *** | 0.06447 |
| Hsp90 interaction score ~ expression maximum<sup>2</sup> | -0.003507 |  | 0.00052 |
| dN/dS ~ expression maximum | -0.0006343 | * | 0.02056 |
| Hsp90 interaction score ~ PPI | -0.35586 | **** | 0.08779 |
| dN/dS ~ PPI | -0.008626 | *** | 0.06256 |

<sup>1</sup>Phylogenetic independent contrasts were used for all regression analyses.
<sup>2</sup>Maximum expression of each kinase was determined across 11 tissues.
p-values: \*0.05, \*\*0.01, \*\*\*0.001, \*\*\*\*0.0001

To further disentangle the contributions of Hsp90 interaction score, gene expression levels, and PPI connectivity to kinase evolutionary rate, we also calculated partial correlations among the contrasts for the four variables. We found that Hsp90 interaction score was positively correlated with kinase evolutionary rate when controlling for both gene expression levels and PPI connectivity (r = 0.19, p = 0.003, Pearson correlation, Table 3). The relative contribution of Hsp90 interaction score and gene expression levels to kinase evolutionary rate was comparable when controlling for the respective other variables; the relative contribution of PPI connectivity was marginal (Table 3). To quantify the combined explanatory power of Hsp90 interaction score, gene expression levels, and PPI connectivity to kinase dN/dS, we modelled dN/dS PIC on the PIC of all three variables. This model explained only a small component of the dN/dS PIC (R^2^ = 0.11).

**Table 3.**
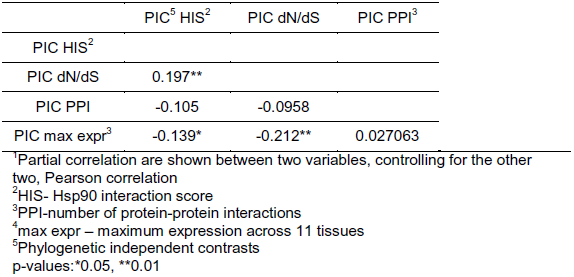
Hsp90 is correlated with dN/dS when controlling for PPI and expression^1^.

|  | PIC<sup>5</sup> HIS<sup>2</sup> | PIC dN/dS | PIC PPI<sup>3</sup> |
| --- | --- | --- | --- |
| PIC HIS<sup>2</sup> |  |  |  |
| PIC dN/dS | 0.197** |  |  |
| PIC PPI | -0.105 | -0.0958 |  |
| PIC max expr<sup>4</sup> | -0.139* | -0.212** | 0.027063 |

<sup>1</sup>Partial correlation are shown between two variables, controlling for the other two, Pearson correlation
<sup>2</sup>HIS- Hsp90 interaction score
<sup>3</sup>PPI-number of protein-protein interactions
<sup>4</sup>max expr – maximum expression across 11 tissues
<sup>5</sup>Phylogenetic independent contrasts
p-values: \*0.05, \*\*0.01

In summary, we found that 1) Hsp90 client status is positively associated with kinase dN/dS, 2) this association appears to be independent of gene expression levels or PPI connectivity, and 3) the strength of association is comparable for Hsp90 client status and gene expression levels. Notably, none of the tested factors alone or in combination explained a large proportion of the observed variation in kinase evolutionary rate.

### Hsp90-associated effects on kinase evolutionary rate are observed across mammals

Thus far, we have analyzed kinase dN/dS values that were calculated from human and mouse, species which diverged ∼75 million years ago (Chinwalla 2002). As mice are known to be a long-branching clade (Wu and Li 1985; Rat-Genome-Sequencing-Project-Consortium 2004), we evaluated whether Hsp90’s effect on kinase dN/dS values was lineage-or time-dependent. To do so, we determined the core set of kinases conserved across all tested mammals and obtained pairwise dN/dS values for the one-to one orthologs of strong human Hsp90 kinase clients and non-clients. In total, 70 kinases were conserved, comprised of 28 non-clients, 20 weak clients, and 22 strong clients. Consistent with the mouse-human comparisons, strong Hsp90 clients evolved faster than non-clients across these species, including several monophyletic lineages such as Primates, Glires, and Laurasitheria (Figure 1b).

Across all comparisons, the average ratio of strong client dN/dS to non-client dN/dS was 1.63±0.25 (standard deviation), indicating that Hsp90 clients undergo relaxed selection compared to non-clients (Figure S1a). Our results suggest that Hsp90 client kinases tend to accumulate non-synonymous variation compared to non-client kinases regardless of species compared and time of divergence.

Several previous studies have demonstrated that Hsp90-dependent standing variation is common in yeast, plant, fly, and fish populations (Rutherford and Lindquist 1998; Queitsch, et al. 2002; Yeyati, et al. 2007; Sangster, Salathia, Lee, et al. 2008; Jarosz and Lindquist 2010). Using genetic or pharmaceutical perturbation of Hsp90, these studies found significantly increased trait heritability upon Hsp90 inhibition, presumably due to the many identified Hsp90-responsive loci. For the vast majority of these Hsp90-dependent loci the identity of the underlying polymorphism (*i.e.* the gene or regulatory region affected) remains unknown and hence evolutionary rate analysis of these loci has yet to be conducted. However, a detailed genetic study found that an Hsp90 client showed increased accumulation of non-synonymous variation compared to its non-client paralog within *A. thaliana* across divergent strains (Lachowiec et al, 2013).

We therefore hypothesized that Hsp90’s effect on kinase evolutionary rate should be detectable among humans. To test this hypothesis, we took advantage of the thousands of sequenced human genomes and examined nucleotide diversity of kinase clients and non-clients as a measure of accumulated genetic variation. Specifically, we assessed nucleotide diversity for kinase clients and non-clients in approximately 6500 individuals (Exome Variant Server), accounting for relatedness and using Hsp90 interaction score contrasts as measure of Hsp90 client status.

Indeed, the PIC of kinase Hsp90 interaction scores and nucleotide diversity were positively correlated (r = 0.157, p = 0.04965, Table S1), suggesting that Hsp90 client kinases harbor greater genetic variation than non-client kinases in humans. This positive correlation suggests that purifying selection is reduced among Hsp90 clients compared to non-clients, resulting in higher levels of slightly deleterious mutations. To determine the type of genetic variation that Hsp90 buffered, we examined nucleotide diversity in non-synonymous and synonymous sites separately. The correlation between the PIC of client score and PIC of nucleotide diversity was stronger when only considering non-synonymous sites (r = 0.23 p = 0.003). In contrast, there was no significant correlation when only considering synonymous sites (r = 0.03, p = 0.69). This result is consistent with the notion that it is Hsp90’s function in protein folding that enables greater dN/dS of its clients rather than the rise of new mutations, epigenetic phenomena, or other, yet unidentified mechanism. To further explore this aspect, we examined whether Hsp90 clients tolerated potentially more deleterious variation across humans, using both Genomic Evolutionary Rate Profiling (GERP) scores (Cooper, et al. 2005) and Polymorphism Phenotyping (PolyPhen) scores (Adzhubei, et al. 2010) as measures of the predicted phenotypic effect of individual single nucleotide variants (SNVs). GERP ascertains the degree of past purifying selection on a site through examination of rejected nucleotide substitutions across homologous sequences (Cooper, et al. 2005), whereas PolyPhen categorizes non-synonymous SNVs by likelihood of damaging protein structure and function (Adzhubei, et al. 2010). Although there was no significant correlation between GERP and Hsp90 interaction score contrasts (r = 0.02, p = 0.7593, Pearson correlation, Table S1), PolyPhen and Hsp90 interaction score contrasts were positively correlated (r = 0.21, p = 0.00987, Pearson correlation, Table S1). To summarize, Hsp90 kinase clients appeared to harbor more genetic variation and more damaging mutations than non-clients across many human individuals.

### Hsp90 client status tends to be gained and maintained

Our observation that Hsp90’s effect on kinase evolutionary rate increases with divergence time suggests that client status is stable over long time periods. This interpretation contrasts with experimental findings that mutating a single amino acid residue can suffice to dramatically alter a protein’s dependence on Hsp90 (Citri, et al. 2006). Similarly, paralogs or otherwise related proteins that differ only by a few amino acids can differ in Hsp90 client status (Citri, et al. 2006; Lachowiec, et al. 2013), suggesting that Hsp90 client status can be highly dynamic. To resolve this apparent contradiction, we analyzed the distribution of Hsp90 client status among gene duplicates in the kinase superfamily. Gene duplicates tended to share Hsp90 client status (Figure 2a, χ^2^-test, p = 1.837e-05) indicating that client status has phylogenetic signal.

**Figure 2.**
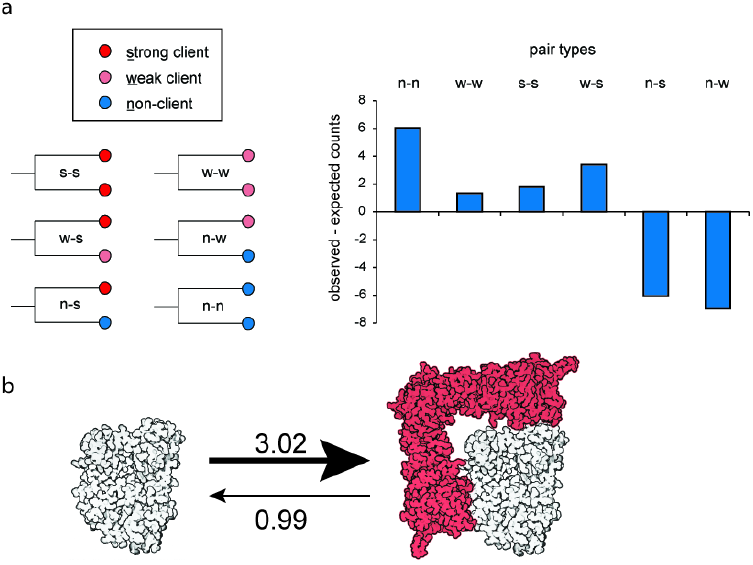
Hsp90 client status dynamics across the kinome. a) Using the three client classes defined by Taipale et al (2012), we categorized pairs of duplicated genes into six classes. We find that gene duplicates share client status (χ^2^-test, p = 1.837*10^−5^) more often than expected by chance. The number of expected pairs was calculated based a random distribution of clients across the tree. b) Kinases (white) are three times more likely to gain Hsp90 client status (red) than lose it based on ML and MCMC estimates of state transition rates using Bayes Traits.

This analysis only considered genes with one paralog, eliminating one-third of the available kinome data. Therefore, we explicitly tested whether there was phylogenetic signal in client status patterns across the entire kinase phylogeny using Pagel’s λ (Pagel 1999). Pagel’s λ is a statistic that tests whether phylogeny correctly predicts the patterns of covariance among species on a given trait. Here, we estimated this statistic to test whether kinase phylogeny predicts the patterns of covariance among kinases with regard to Hsp90 client status. Assuming that kinases were either Hsp90 clients or non-clients, we compared a model in which λ was computed using the known kinase phylogeny (Manning, et al. 2002) (λ = 0.74) to a model assuming no phylogenetic signal (λ = 0, star phylogeny). The former model including phylogenetic signal was a better fit for the observed pattern of client status across the kinase phylogeny than the model without phylogenetic signal (2δ = 8.94, df = 1, p = 0.00279). We conclude that client status is not randomly distributed, and that more closely related kinases tend to share client status across the whole kinase superfamily.

Having established that client status tends to be shared among related kinases, we next explored the evolutionary paths by which client status is acquired and changes through time. To do so, we estimated the transition rates between client and non-client states along the kinase phylogeny, using maximum likelihood (ML) implemented in the software BayesTraits (Pagel et al. 2004; Pagel 1999). We found that the rate of the transition from non-client to client along the kinase phylogeny was 3.02, compared to the reverse rate of 0.99. Restricting these rates to equal values significantly worsened the model fit (2δ = 8.066, df = 1, p = 0.0045), hence we conclude that kinases are more likely to become Hsp90 clients than to lose client status (Figure 2b). To examine whether these ML-derived rates were robust, we also used the Markov Chain Monte Carlo (MCMC) analysis implemented in BayesTraits to estimate the rates of gain and loss of client status. The rates of transition for non-client to client (3.11) and client to non-client (0.98) indicated that rate estimates were robust (Figure 2b). The tendency of non-client kinases to become clients and of client kinases to remain clients agrees with previous suggestions that kinase dependence on Hsp90 may be ‘addictive’ and may contribute to the greatly increased sensitivity of cancer cells to Hsp90 inhibitors (Workman, et al. 2007).

If protein kinases indeed become ‘addicted’ to being Hsp90 clients, one may expect the ancestral kinase to be a non-client. To infer the client status of the ancestral kinase, we compared a tree with a non-client kinase root to a tree with a client kinase root using MCMC. We found strong support for the tree with a non-client root (Bayes factor ∼9.1). Both Hsp90 and kinases were present in the eukaryotic common ancestor (Manning G 2009; Bogumil, et al. 2014); however, our results would suggest that they likely did not interact. Although, Hsp90 client predictions and confirmed clients in the prokaryote *E. coli* include kinases (Press, et al. 2013), these prokaryotic protein kinases belong to a different kinase family and differ in structure (Manning G 2009). Taken together, our results are consistent with a scenario in which early kinases evolved independently of Hsp90 with subsequent multiple independent gains of Hsp90 client status, increasing Hsp90’s effect on kinase evolution and adding to the overall chaperone dependence of kinases for their function.

## Discussion

Although previous studies showed that Hsp90-dependent standing variation is common in natural populations (Rutherford and Lindquist 1998; Queitsch, et al. 2002; Sangster, Salathia, Lee, et al. 2008; Jarosz and Lindquist 2010; Rohner, et al. 2013), the chaperone’s impact on protein evolution, especially in comparison to established factors such as gene expression, has remained unknown. Here, we present evidence for a direct role of Hsp90 in kinase evolution and compare its impact to gene expression levels and PPI connectivity.

We find that interaction with Hsp90 is associated with higher dN/dS for kinases and that Hsp90 client status appears to contribute to kinase evolutionary rate independently of gene expression and PPI connectivity. Each of these factors contributed to a similar degree to kinase evolutionary rate; combined they explained only about 10% of kinase dN/dS variation. Adding additional variables may decrease the amount of unexplained variation, but many variables are known to co-vary. For example, Bloom and Adami (2003) have argued that PPI connectivity is confounded by protein abundance, with highly abundant proteins engaging in a larger number of interactions (Bloom and Adami 2003). Protein abundance arises as a combination of gene expression levels, and rates of translation and protein degradation. Others have argued that the dominant role of gene expression in driving evolutionary rate is at least in part due to selection for translational robustness, which encompasses selection for increased translational accuracy by optimizing codon usage, and selection to increase the number of proteins that fold properly despite mistranslation (Drummond, et al. 2005). Previously, optimized codon usage, as measured by codon adaptation index, was observed for stochastic clients of GroEL/ES (Warnecke and Hurst 2010). However, we did not observe an association between CAI and Hsp90 client score (p = 0.985, R^2^=2.015*10^−6^), possibly reflecting that Hsp90 does not act as a co-translational chaperone (Taipale, et al. 2010) like GroEL/ES (Ying, et al. 2006). Further, large-scale studies of translation efficiency and its correlation with CAI in human have yielded contradictory results, with some studies failing to find significant correlations (Chamary, et al. 2006; Waldman, et al. 2010). We also note that the kinase Hsp90 clients analyzed here are generally less abundant in steady-state protein levels (Taipale, et al. 2012), presumably due to their enhanced structural lability. Nevertheless, by using partial contrasts, we find evidence for a contribution of Hsp90 to kinase dN/dS independent of expression level and PPI degree.

As suggested by prior studies of Hsp90-dependent variation within diverse populations of plants, yeast, fly and fish (Rutherford and Lindquist 1998; Queitsch, et al. 2002; Yeyati, et al. 2007; Sangster, Salathia, Undurraga, et al. 2008; Jarosz and Lindquist 2010), we found support for Hsp90-dependent genetic variation across humans. Both nucleotide diversity and PolyPhen scores were significantly correlated with kinase Hsp90 interaction scores; yet effect sizes were modest. No significant correlations were observed for GERP scores. The GERP score is a position-specific estimate of evolutionary constraint using maximum likelihood evolutionary rate estimation (Cooper, et al. 2005). High evolutionary constraint is often interpreted as functional relevance. As we do not expect Hsp90-dependent variation to reside in highly constrained sites, it is not surprising that GERP scores for Hsp90 kinase clients and non-clients did not differ significantly.

In agreement with the modest signal of increased nucleotide diversity within human, we observed an increased evolutionary rate for Hsp90 client kinases across divergent mammalian lineages, consistent with Hsp90 allowing for greater accumulation of non-synonymous changes in the genes encoding its clients. In fact, Hsp90’s effect on kinase evolutionary rate increases when considering species that are more distantly diverged. The greater accumulation of non-synonymous changes may allow client kinases to explore a wider sequence space and potentially acquire novel functions at a faster rate.

Taipale et al (2012) also determined Hsp90 interaction scores for a large number of transcription factors and E3 ligases. In contrast to our findings for client and non-client kinases, transcription factor and E3 ligase Hsp90 interaction scores were not significantly associated with their evolutionary rates (Text S1, Figure S2). Unlike the kinases analyzed, the transcription factors are not monophyletic (Vaquerizas, et al. 2009) and hence may differ more in other factors influencing evolutionary rate, such as protein structure and stability. As for the E3 ligases, others have suggested that Hsp90 works in concert with E3 ligases to promote proteasome-dependent degradation rather than chaperoning these enzymes (Murata, et al. 2001; McClellan, et al. 2005; Morishima, et al. 2008; Ehrlich, et al. 2009; Taipale, et al. 2010). Hsp90 works in concert with many other proteins that enable and modify its function and generate client specificity, such as diverse Hsp70s, co-chaperones, and immunophilins (Taipale, et al. 2010). Although these “collaborating” proteins physically interact with Hsp90, they do so in a sequence-or domain-specific manner, and Hsp90 does not facilitate their folding (Murata, et al. 2001; McClellan, et al. 2005; Morishima, et al. 2008; Ehrlich, et al. 2009; Taipale, et al. 2010); hence they are unlikely to experience relaxed selection due to this interaction. Following this line of reasoning, we previously excluded known Hsp90 co-chaperones and Hsp70s from evolutionary rate analyses (Lachowiec, et al. 2013).

Previously, we found that Hsp90 clients in plants and yeast showed significantly greater evolutionary rates than their non-client paralogs (Lachowiec et al. 2013). The kinome data set only contained five pairs of strong kinase clients and non-clients. Although, in four of these pairs the strong kinase client showed greater dN/dS than its non-client paralog, this difference was not significant, presumably due to small sample size (n=5, 95% confidence interval 0.6-12.6, p=0.125, one-sample Wilcoxon test, testing the deviation from the expected ratio of client/non-client dN/dS of 1). We did, however, detect a significant trend for duplicate genes such that paralog pairs that contained at least one strong Hsp90 client showed significantly greater divergence than pairs that did not (Figure S3). We speculate that interaction with Hsp90 allows clients to tolerate slightly deleterious, but non-lethal mutations without losing function, thereby facilitating the emergence of novel functions over time (Lachowiec et al. 2013).

Beyond paralog pairs, we further explored the relationship of Hsp90 client status and kinase evolutionary rate among kinase families. The effect of Hsp90 client status on dN/dS was consistent across all kinase groups (Figure S4a) with greater dN/dS observed for Hsp90 client kinases compared to non-client kinases. The TK and TKL families were enriched for strong Hsp90 clients (Figure S4b), and possibly in part due to this enrichment, both families showed the highest evolutionary rates among the tested kinase families (Figure S4c). TK and TKL kinase families are evolutionarily young and have been implicated in the rise of multicellularity (Lim and Pawson 2010). In contrast, of all families, the CAMK family was most depleted for strong Hsp90 clients. Unlike in all other kinase families, the few strong CAMK clients did not evolve faster than the weak clients (Figure S5). As almost a third of the CAMK family genes are pseudogenes (not included in this analysis), contrasting with only about one-fifth of kinase pseudogenes overall (Manning, et al. 2002), we speculate that the “missing” strong CAMK clients have become pseudogenes.

These family-specific observations and our finding that kinases tend to acquire and maintain Hsp90 client status prompt our speculation that Hsp90 may play a complex role in the birth and death of kinases. The most common outcome after gene duplication is pseudogenization of one copy (Nei and Roychoudhury 1973). Acquiring Hsp90 client status likely leads to instant sub-functionalization due to the temperature sensitivity of clients and may facilitate gene copy maintenance (Lachowiec, et al. 2013). The observed greater accumulation of non-synonymous variation in genes encoding Hsp90 client may also facilitate neo-functionalization. At the same time, the greater accumulation of more harmful variation may predispose genes encoding clients to the pseudogene fate. In other words, acquiring Hsp90 client status may be akin to a delayed death sentence on a long evolutionary time scale.

In fact, the suggested “Hsp90 addiction” of kinases (Workman, et al. 2007), especially in cancer cells with their mutated oncogenic kinases, is reminiscent of a recent argument by Fernandez and Lynch (Fernández and Lynch 2011). These authors attributed the increasing complexity of protein interaction networks from bacteria to human to compensation for the decreased stability of proteins over evolutionary time. They posited that through drift proteins would be exposed to destabilizing mutations and hence become susceptible to aggregation and malfunction. Interaction in homo-and hetero protein complexes will then compensate for the stability deficits of proteins in higher organisms (Fernández and Lynch 2011). Of course, chaperones also prevent aggregation and stabilize proteins (Taipale, et al. 2010). The small evolutionary snapshot of the kinome with its tendency to acquire and maintain Hsp90 client status fits well within this framework of thought.

## Methods

### Data sources and estimating contributions to dN/dS

Hsp90 interaction scores, client category, and number of connections in the protein-protein interaction (PPI) network were obtained from (Taipale, et al. 2012). For gene expression levels in human, RNA-seq counts normalized for read depth (RPKM, Reads per kilo-base per million) across eleven normal human tissues from (Castle, et al. 2010) were downloaded (http://medicalgenomics.org/rna_seq_atlas/ [September, 2012]). The average expression was taken per gene (across all splice forms) and the maximum and mean expression across all tissues was calculated. The DNA sequence of the longest splice form for each kinase was obtained from Ensembl release 70, (http://www.ensembl.org [March, 2013]). Codon usage bias for humans was obtained from http://www.kazusa.or.jp/codon/ (Nakamura, et al. 2000). The codon adaptation index for each kinase was calculated using the first 50 and last 20 codons (Warnecke and Hurst 2010) using E-CAI (http://genomes.urv.cat/CAIcal/E-CAI/) (Puigbo, et al. 2008). Only the 210 kinases that overlapped between the Hsp90 kinome analysis (Taipale, et al. 2012) and the eukaryotic kinase tree (Manning, et al. 2002) and that had values for expression (Castle, et al. 2010), dN/dS, and PPI connectivity were used in the downstream analyses. Since the phylogenetic relatedness of the kinases may influence these contributions, we also considered each variable in a phylogenetic context using phylogenetically independent contrasts (PIC). Phylogenetic contrasts for each variable were estimated in R using *pic* in the package ape (Paradis, et al. 2004) with the kinome tree from (Manning, et al. 2002) (http://kinase.com/human/kinome/groups/ePK.ph). We conducted linear regression analyses with the intercept set to 0 (Garland et al., 1992) since the order of subtraction to calculate the PIC was arbitrary and calculated the association between each pair of variables. Partial correlations among the variables were calculated with R using *pcor* in the package ppcor (http://cran.r-project.org/web/packages/ppcor/index.html).

### dN/dS comparisons across species

For each kinase, orthologs for the human kinases and their respective dN/dS values were identified using Ensembl release 70, (http://www.ensembl.org [March, 2013]) in the following species: *Pan troglodytes*, *Pongo abelii*, *Nomascus leucogenys*, *Macaca mulatta*, *Callithrix jacchus*, *Mus musculus*, *Rattus norvegicus*, *Ictidomys tridecmlineatus*, *Oryctolagus caniculis*, *Canis familiaris*, *Bos taurus, Sus scrofa*, and *Ailuropoda melanoleuca*. If multiple orthologs were identified, the kinase was removed from the analysis. Time to common ancestor was obtained from compiled studies curated at TimeTree.org (Hedges et al. 2006).

### Hsp90 client divergence timing among humans

We acquired human exome sequences from the Exome Variant Server (Exome Variant Server, NHLBI GO Exome Sequencing Project (ESP), Seattle, WA (http://evs.gs.washington.edu/EVS/ [November, 2012]) using an in-house Perl script.

In addition to the filters from the ESP Server, we incorporated the filters used in (Tennessen, et al. 2012): all SNVs had to have a quality score above 20, allele balancing above 65%, read depth between 10 and 1000x to account for possible copy variation, and we excluded genotype qualities of zero.

To calculate nucleotide diversity, we used

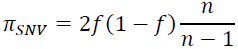

where *f* is the frequency of the major allele and *n* is the number of haploid genomes (Hernandez et al. 2011). For each gene *π_SNV_* was summed for each nucleotide position and normalized for gene length. For gene length we used the length of the longest CDS reported by Ensembl release 69.

We extracted GERP scores for each SNV from ESP and calculated the average GERP score for each kinase gene and normalized the GERP counts by the length of the gene (CDS sequence). PolyPhen scores were also obtained from ESP.

### Examining client status dynamics and estimating transition rates between client and non-client states

We estimated the phylogenetic signal of client status across the kinome, Pagel’s λ, using the function *fitDiscrete* in the package geiger (http://cran.r-project.org/web/packages/geiger/index.html). We modeled a kinase tree with all branches leading to tips of equal length (λ = 0) eliminating the phylogenetic signal. We then compared this model to a model with the true kinase phylogeny (i.e. an optimized λ) using likelihood-ratio tests, where the likelihood ratio approximates a χ^2^ distribution with one degree of freedom (Harmon, et al. 2008; Motani and Schmitz 2011).

We used BayesTraits v2 (Pagel 1994)(http://www.evolution.rdg.ac.uk/BayesTraits.html) to estimate transition rates along the kinase tree using a MultiState model of evolution. We used the human kinase tree and the client status of each kinase, coding both weak and strong clients (according to Taipale et al, 2012) as ‘A’ and non-clients as ‘B’. We then used maximum likelihood with the parameter rate set to 2, representing the two transition rates 1) client to non-client and 2) non-client to client. We tested various hypotheses about transition rate parameters by restricting transition rates: 1) rate of gain of client status to 0, 2) rate of loss of client status to 0, and 3) equal gain and loss rates. We compared the log(likelihood) of the restricted models to one another and to the log(likelihood) of the unrestricted models. For these comparisons we used likelihood-ratio tests, where the likelihood ratio follows a χ^2^ distribution with degrees of freedom equal to the number of restricted parameters. We repeated the analysis using Markov Chain Monte Carlo (MCMC) implemented in BayesTraits v2. We had appropriate levels of acceptance with the option rateDev set to 2. We let the chain run for 100,000 iterations with a uniform prior distribution between 0 and 100. We examined the transition rates after a 20,000 iteration burn-in period.

To determine the client status of the root kinase, we used MCMC implemented in BayesTraits v2. We used the kinase tree and client status coding described above. We fossilized the root as either A or B for the whole tree. We found appropriate levels of acceptance with the option rateDev set to 1. We ran the chain for 10 million iterations, and compared the two different client states of the root by calculating the Bayes Factor based on the harmonic means (twice the difference between the two harmonic means of the model likelihood) (Pagel, et al. 2004).

### Examining non-kinase Hsp90 interactors

We removed all genes that were found in more than one functional category: E3 ligases, transcription factor, or kinase. The client status as defined in Taipale et al (2012) was used for each gene, with E3 ligases and transcription factors (TFs) categorized as “not significant interactor”, removed from further analyses. We used pairwise dN/dS values with mouse as an outgroup to examine the dN/dS among the Hsp90 interactors and non-interactors. Neither TFs nor E3 ligases are monophyletic. We subdivided the TFs into phylogenetically related families (Vaquerizas, et al. 2009) and compared the dN/dS between Hsp90 interactors and non-interactors within each family and in aggregate. Subdividing E3 ligases into phylogenetic groups based on sequence similarities has not been previously conducted, so we subdivided E3 ligases based on domain presence (Kelch, WD40 from (Taipale, et al. 2012) or RING, U box, HECT, F box, SOCS box, BTB, DDB1-like, ZnF A20 from (Li, et al. 2008)) and compared the dN/dS between Hsp90 interactors and non-interactors within each group. Because no differences in dN/dS were found between Hsp90 interactors and non-interactors within groups defined by individual domains, we also clustered E3 ligases based on presence or absence of many domains simultaneously. E3 domains were identified using PFAM version 26. Clustering was completed using hierarchical clustering in R with the function *heatmap*.

## Acknowledgments

We thank Maximilian Press, Joe Felsenstein, and Mikko Taipale for helpful discussions. We thank the NHLBI GO Exome Sequencing Project and its ongoing studies which produced and provided exome variant calls for comparison: the Lung GO Sequencing Project (HL-102923), the WHI Sequencing Project (HL-102924), the Broad GO Sequencing Project (HL-102925), the Seattle GO Sequencing Project (HL-102926) and the Heart GO Sequencing Project (HL-103010). This work was supported by the National Science Foundation (DGE-1256082 to JL), the National Institutes of Health (DP2OD008371 to CQ, DP2AT00780201 to EB), and the Alfred P Sloan Foundation (to EB).

